# Ventral tegmental area GABA neurons integrate positive and negative valence

**DOI:** 10.1101/2024.12.07.627330

**Authors:** Margaret E. Stelzner, Amy R. Wolff, Benjamin T. Saunders

## Abstract

The ventral tegmental area (VTA) is classically linked to reward learning and reinforcement via the actions of dopamine neurons. Intermingled VTA GABA neurons are positioned to regulate dopaminergic systems, but we lack detailed insight into how this population contributes to conditioned motivation across learning contexts. Here, recording population calcium activity in VTA dopamine and GABA neurons during different conditioning paradigms, we found that, while dopamine neurons were selectively responsive only to appetitive stimuli, GABA neurons actively encoded both appetitive and aversive cues and outcomes. GABA neurons also signalled errors in violation of expectations for associations of either valence. Critically, we found that GABA neurons selectively integrated events when both valences were simultaneously present, reflecting motivational state under cost. In a motivational conflict task, where rats must weigh decisions to avoid shock or seek reward, VTA GABA neuron activity scaled with escalating cost, and was predictive of reward seeking motivation. Optogenetic inhibition of VTA GABA neurons disrupted reward seeking specifically under conditions of motivational conflict. Together, our data show that VTA GABA neurons reflect a broader learning signal, compared to dopamine neurons, one that is particularly important for directing appropriate behavioral responses in complex, multivalent environments. Our results reveal new insights into the functional landscape of the VTA, where distinct populations of neurons act in parallel to signal different aspects of motivation under various behavioral conditions.

## Introduction

The Ventral tegmental area (VTA) is centrally involved in learning and reinforcement (Berke, 2018; Cox & Witten, 2019; Howe & Dombeck, 2016; Schultz, 2006; Sharpe et al., 2017; Tsai et al., 2009). Classically, VTA dopamine neurons encode cue-outcome associations with increased activity to positively valenced events and decreased activity to negatively valenced events (Cohen et al., 2012; Day et al., 2007; Keiflin et al., 2019; Keiflin & Janak, 2015; Morales & Margolis, 2017; Saunders et al., 2018; Schultz et al., 1997; Schultz, 2016; Steinberg et al., 2013). The VTA, however, is a heterogeneous region and also contains a substantial population of GABAergic neurons that are less understood (Breton et al., 2019; Nair-Roberts et al., 2008; Oriol et al., 2024; Swanson, 1982).

VTA GABA neurons have been seen as an “anti-reward” signal because of their ability to locally inhibit dopamine neurons (Corre et al., 2018; Tan et al., 2012; van Zessen et al., 2012). Growing evidence over the past decade has expanded this notion, demonstrating a role for VTA GABA neurons in behavioral flexibility, motivation and reward, and memory (Al-Hasani et al., 2021; Bouarab et al., 2019; Cohen et al., 2012; Elum et al., 2024; Eshel et al., 2015; Glykos & Fujisawa, 2024; Hughes et al., 2019; Kim et al., 2010; Lefner & Moghaddam, 2025; Ostroumov & Dani, 2018; Zhou et al., 2022), with inconsistent evidence of the active role of these neurons in appetitive versus aversive contexts. VTA GABA neuron activity has primarily been considered in relation to dopamine computations, where they are thought to contribute to reward prediction error by inhibiting dopamine neurons during expectation and omission of reward (Cohen et al., 2012; Eshel et al., 2015). Critically, there remains a dearth of direct comparisons between VTA dopamine and GABA neuron encoding as parallel processors in multi-valent learning contexts, where unique motivational resources may be marshalled to promote decision-making under conflict. This presents an important gap to developing more complete models of VTA function that extend beyond simple reward learning (Bromberg-Martin et al., 2010; Gershman et al., 2024; Lee et al., 2024; Schultz, 2013).

To address these questions, we selectively targeted dopamine or GABA neurons of the VTA with GCaMP and recorded population-level calcium fluorescence as a proxy for neural activity in freely moving rats engaged in various learning paradigms. Our results add new insight to emerging perspectives that broaden the computational and functional landscape for the VTA, where distinct populations of neurons act in parallel, rather than opposition, to signal different aspects of motivation under dynamic behavioral conditions. We find that, while dopamine neurons are selectively responsive to appetitive, but not aversive stimuli, GABA neurons actively encode both appetitive and aversive cues and outcomes. Further, GABA neurons signal errors in violation of expectations for both appetitive and aversive associations, and integrate events when both valences are simultaneously present. In a motivational conflict task, where rats must weigh decisions to avoid shock or seek reward, we found that VTA GABA neuron activity scaled with escalating cost, and was predictive of reward-seeking motivation when cost was present. Brief optogenetic inhibition of VTA GABA neurons disrupted reward seeking specifically under conditions of motivational conflict. Together our data suggest that VTA GABA neurons engage a broader learning mechanism, compared to dopamine neurons, one that is important for directing appropriate behavioral responses in complex, multivalent environments.

## Results

### Conditioned and unconditioned appetitive events are similarly encoded by VTA dopamine and GABA neurons

Expression of the calcium indicator GCaMP8f was targeted to dopamine or GABA neurons in the VTA, using a combination of transgenic rats and promoter-driven viral tools (Fig. 1A-C; Supplemental Fig. 1). We first trained rats on a Pavlovian reward conditioning paradigm in which a conditioned stimulus (CS+) predicted reward delivery (liquid Ensure) and a distinct neutral stimulus (CS-) had no paired outcome (Fig. 1D), across 15 sessions. Rats reliably distinguished between the stimuli, with growing discrimination across Early (day 1), Middle (day 8 or 9) and Late (day 13 or 15) training phases, coming to selectively approach the reward port in response to the CS+ (Fig. 1E; two-way repeated measures ANOVA, cue-type main effect, F(1,36)=20.94, p<0.0001; session by cue interaction F(2,66)=12.92, p<0.0001), and doing so with faster latencies (Fig. 1F: one-way RM ANOVA, session main effect, F(1.7,31.1)=95.69, p<.0001). Using video tracking of the rat’s position in the chamber, we saw that approaches to the reward port were initiated rapidly after CS+ onset (Fig. 1G). We found no behavioral differences across sex or neuronal recording groups (Supplemental Fig. 2). Across all subjects, final fiber placements were centered over the middle of the VTA (mean: +.75 mm lateral from midline; range: +.5 mm to +1.1mm lateral) and we found no significant correlations between recording location and neural activity. Robust CS+-evoked activity emerged in both dopamine and GABA neurons as learning progressed, with similar dynamics (Fig. 1H, P). This activity also consistently increased to reward consumption throughout conditioning in both neural subtypes (Fig. 1J, R). Dopamine (Fig. 1H-O) neurons strongly discriminated CS+ from CS-presentations with increasing CS+ preference across training, as measured by area under the curve (AUC; Fig 1L; session by cue interaction, F(1.77,17.67)=7.19, p=.0065; main effect of cue, F(1,10)=10.24, p=.0095) and peak signal measures (Fig. 1M, session by cue interaction, F(1.95,19.45)=8.587, p=.0023; main effect of cue, F(1,10)=5.016, p=.049). Dopamine neuron activity strongly discriminated rewarded versus unrewarded port entries, consistently across training, for AUC (Fig. 1N: session by response type interaction, F(1.47,14.7)=.1531, p=.795; effect of response type, F(1,10)=41.12, p<.0001) and peak signal measures (Fig 1O; session by response type interaction, F(1.47,14.65)=.134, p=.812; effect of response type, F(1,10)=58.51, p<.0001). Similar to dopamine, GABA neuron (Fig. 1P-W) signals strongly discriminated CS+ from CS-presentations for AUC (Fig 1T; session by cue interaction, F(1.88,13.19)=16.89, p=.0003; main effect of cue, F(1,7)=58.66, p=.0001) and peak measures (Fig 1U; session by cue interaction, F(1.58,11.1)=12.29, p=.0023; main effect of cue, F(1,7)=79.37, p<.0001). GABA neurons also responded strongly to reward, discriminating rewarded from unrewarded port entries for AUC (Fig. 1V; interaction, F(1.71,11.96)=.2605, p=.855; effect of response type, F(1,76)=5.606, p=.049) and peak (Fig. 1W; interaction, F(1.44,10.1)=.1.589, p=.246; effect of response type, F(1,7)=8.432, p=.023) measures.

**Fig 1.**
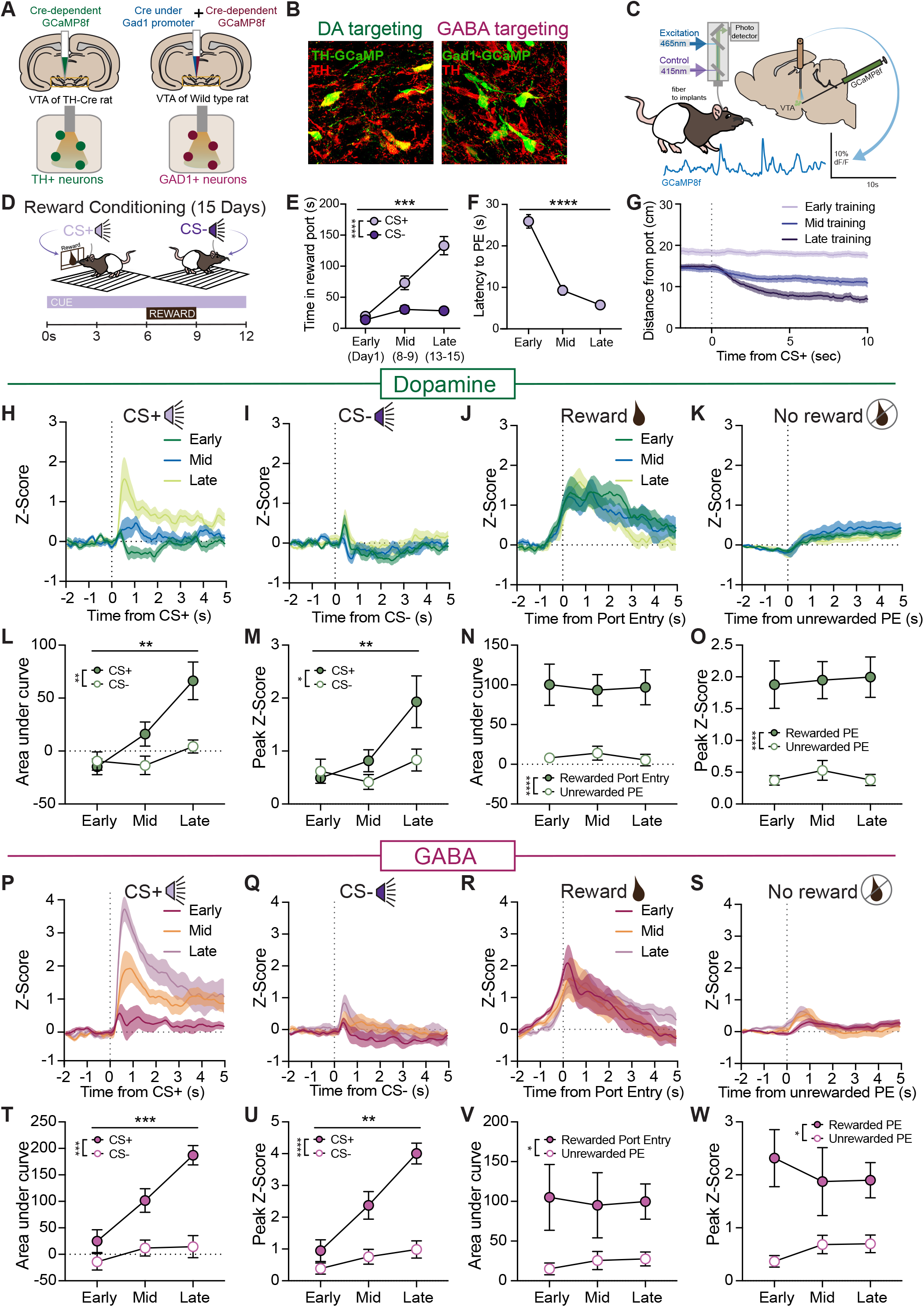
Conditioned and unconditioned appetitive events are similarly encoded by VTA DA and GABA neurons. **A)** Approach for cell-type specific targeting with dopamine neurons targeted through a TH-cre transgenic line and GABA neurons targeted with a viral delivery of cre to GAD1+ neurons. **B)** Examples of viral expression in the VTA. **C)** Schematics of fiber photometry approach, and **D)** experimental design. Rats received cue (CS+) + reward (Ensure) pairings and a distinct neutral cue (CS-) that did not have a paired outcome, while recording VTA GCaMP signals with fiber photometry. **E)** Rats (n=19; 8F, 11M) learned to discriminate the two cues as measured by time spent in reward port during conditioned stimuli presentations across Early (day 1), Middle (day 8 or 9) and Late (day 13 or 15) training phases. **F)** Port entry latency following CS+ presentations decreased across training phases. **G)** As rats learned, they initiated approaches to the reward port rapidly after CS+ onset. **H-O)** Calcium recordings from VTA dopamine neurons (n=11; 4F, 7M). **H)** Dopamine neurons developed robust activity in response to the CS+ cue, **I)** while showing minimal responses to the CS- cue. **J)** Reward consumption evoked consistent dopamine neuron activity across training, **K)** while unrewarded port entries (PEs) were associated with minimal dopamine activity. **L)** Dopamine neuron activity strongly discriminated CS+/CS- cues as training progressed, as measured by AUC and **M)** peak measures of the GCaMP signal. **N)** Dopamine neuron activity strongly discriminated rewarded versus unrewarded port entries consistently across training for both AUC and **O)** peak signal measures. **P-W)** Calcium recordings from VTA GABA neurons (n=8; 4F, 4M). **P)** GABA neurons developed robust activity to the CS+ cue, **Q)** while showing minimal responses to the CS- cue. **R)** Reward consumption evoked consistent GABA neuron activity across training, **S)** while unrewarded port entries were associated with minimal GABA activity. **T)** GABA neuron activity strongly discriminated CS+/CS- cues as training progressed, as measured by AUC and **U)** peak measures of the GCaMP signal. **V)** GABA neuron activity also strongly discriminated rewarded versus unrewarded port entries consistently across training for both AUC and **W)** peak signal measures. Data reflect subject means +/- SEM. *p< 0.05, **p≤ 0.01, ***p≤ 0.001, ****p≤ 0.0001.

**Fig 2.**
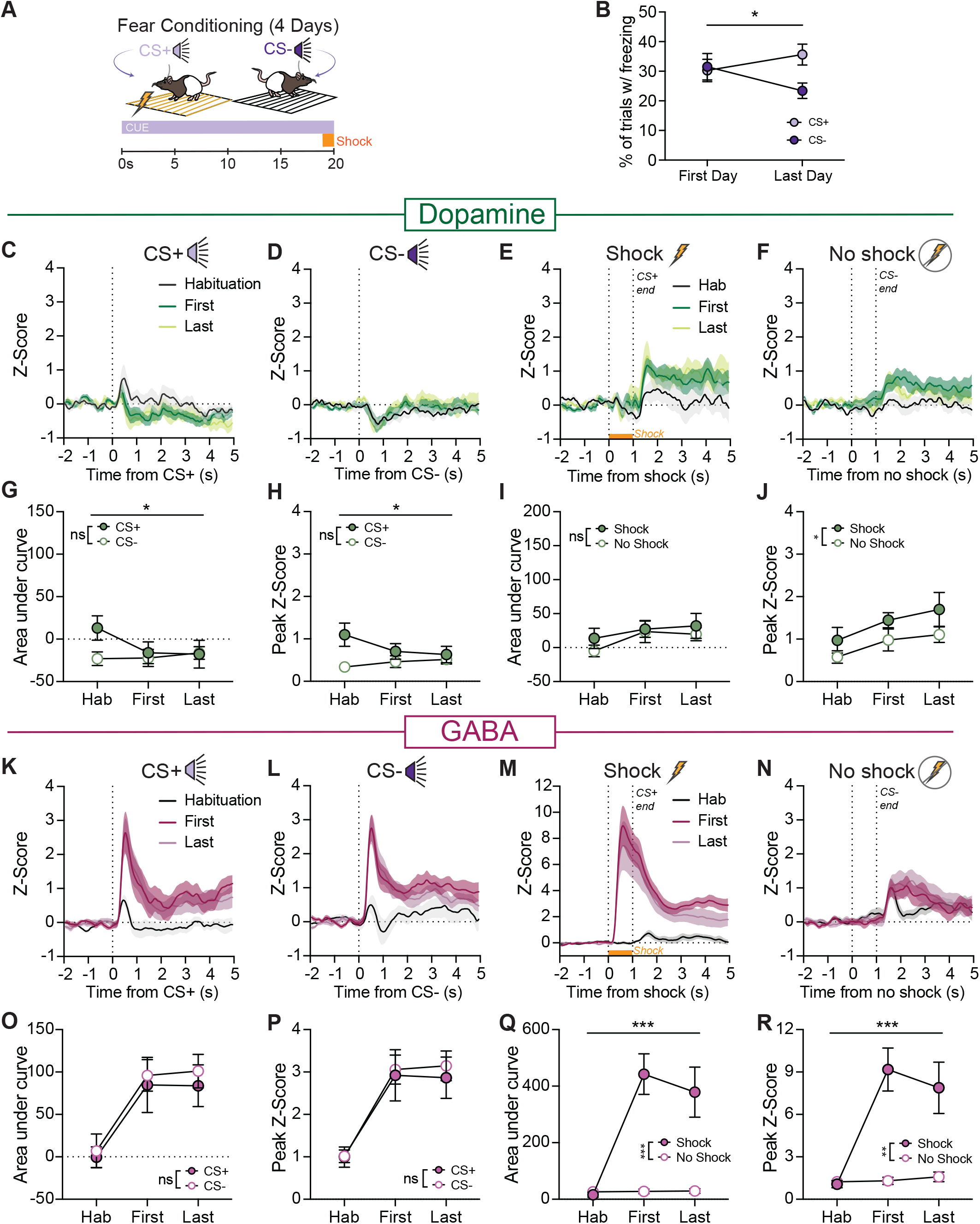
Conditioned and unconditioned aversive events are strongly encoded by VTA GABA, but not VTA DA neurons. **A)** Schematic of fear conditioning design. After a cue habituation session, rats received cue (CS+) + shock (0.4mA, 1s) pairings and a distinct cue (CS-) with no paired outcome, while we recorded GCaMP signals with fiber photometry for 3 sessions. **B)** By the end of conditioning, rats learned to discriminate the CS+/CS- cues, as measured by % of trials with a freezing response. **C-J)** Calcium recordings from VTA dopamine neurons (n=9). **C)** Dopamine neurons showed minimal responses to the onset of the CS+ and **D)** CS- cues across cue habituation and fear conditioning phases. **E)** There was no dopamine neuron activity in response to shock itself, and small increases to the offset of the CS+/shock and **F)** offset of the CS- cue. **G)** Across conditioning, dopamine neuron activity did not discriminate CS+/CS- cues as measured by AUC and **H)** peak measures of the GCaMP signal. **I)** Dopamine neuron activity did not discriminate between the shock period and “no shock” periods corresponding to the end of the CS+ and CS- cues, based on AUC. **J)** Dopamine neuron activity at shock offset and CS- offset did increase slightly across conditioning, and the peak of this shock offset signal was larger than the CS- offset peak. **K-R)** Calcium recordings from VTA GABA neurons (n=13). **K)** GABA neurons developed strong responses to the onset of the CS+ and the **L)** CS- cues between the habituation and conditioning phases. **M)** Shock evoked large, consistent GABA neuron responses, and **N)** the offset of the CS- cue evoked small increases in activity across conditioning. **O)** Strong GABA neuron activity developed to both cues as conditioning progressed, and did not discriminate CS+/CS- according to AUC and **P)** peak signal measures. **Q)** Robust GABA activity was evoked by shock, relative to the comparable period of time at the end of the CS- cue, based on AUC and **R)** peak measures. Data reflect subject means +/- SEM. *p< 0.05, **p≤ 0.01, ***p≤ 0.001, ****p≤ 0.0001.

Together, these results show that, at the population level, GABA neurons are activated by appetitive stimuli with a similar pattern and time course as dopamine neurons. Broadly, the similarity in appetitive event encoding between dopamine and GABA neurons supports the notion that they can exhibit parallel, as opposed to merely oppositional, roles in reinforcement learning.

### Conditioned and unconditioned aversive events are strongly encoded by VTA GABA, but not VTA dopamine neurons

In some prevailing frameworks, dopamine neurons are inhibited by aversive stimuli, potentially via activity of local GABA neurons (Cohen et al., 2012; Schultz, 2016; Tan et al., 2012). Other findings, however, demonstrate that VTA dopamine neurons can play an active role in Pavlovian threat and punishment encoding (Brischoux et al., 2009; Cai et al., 2020; Jo et al., 2018; Park & Moghaddam, 2017). It therefore remains unclear how these populations directly compare in their engagement with aversive learning.

To explore this, we next trained rats on a fear conditioning paradigm, with presentations of a conditioned stimulus predicting a moderate footshock (CS+) and a neutral stimulus (CS-) (Fig. 2A), across 3 sessions, while recording dopamine or GABA neurons. Rats learned to distinguish the two cues as measured by their freezing behavior (Fig. 2B; two-way ANOVA, cue-type x session interaction F(1,28) = 4.943, p=0.0344). In contrast to their patterns of activity in reward conditioning, VTA dopamine neurons (Fig. 2C-J) showed minimal response to the onset of the shock CS+ and did not discriminate the CS+/-for AUC (Fig. 2C,G; phase by cue type interaction, F(1.67,13.33)=5.55, p=.022; main effect of cue type, F(1,8)=2.07, p=.189) or peak measures (Fig. 2D,H; session by cue interaction, F(1.7,13.7)=5.32, p=.023; main effect of cue, F(1,8)=5.03, p=.055). Dopamine neurons also did not respond to the shock itself, and did not discriminate between the shock period and “no shock” periods corresponding to the end of the CS+ and CS-cues, based on AUC (Fig 2E,I; effect of shock condition, F(1,8)=1.32, p=.283). A small increase in dopamine neuron activity was evident at the point of shock offset, which was larger than for the offset of the CS-based on the peak measure (Fig. 2E,F,J; effect of phase, F(1.65,13.2)=6.76, p=.012); effect of shock condition, F(1,8)=7.77, p=.024).

GABA neurons (Fig. 2K-R) exhibited a wholly different pattern, rapidly signaling the onset of the cues during conditioning, with no significant discrimination between the CS+ and CS-for AUC (Fig. 2O; effect of phase, F(1.81,21.68)=19.45, p<.0001); effect of cue type, F(1,12)=1.25, p=.285) and peak (Fig. 2P; effect of phase, F(1.88,22.55)=20.70, p<.0001); effect of cue type, F(1,12)=.229, p=.641) measures. GABA neurons were also strongly engaged by shock itself, relative to the comparable no shock period of time at the end of the CS-, for AUC (Fig. 2Q; phase by shock condition interaction, F(1.85,19.39)=14.17, p<.0001), effect of shock condition, F(1,12)=27.35, p<.0001) and peak measures (Fig. 2R; phase by shock condition interaction, F(1.84,22.1)=11.05, p<.0001), effect of shock condition, F(1,12)=18.33, p<.0001). These data help clarify ongoing research indicating that VTA GABA neurons, relative to dopamine neurons, are strongly responsive to aversive conditioning, implying a broader role in associative learning.

### VTA GABA neurons differentially track appetitive and aversive expectation violations

Dopamine neuron activity in response to expectation violations has been categorized as a reward prediction error (Schultz et al., 1997). Classically, dopamine neurons increase activity when an outcome is better than expected and decrease their activity when an outcome is worse than expected. GABA neurons have been shown to be an important modulator of this dopaminergic prediction signal by encoding expectation of reward, via ramping signals (Cohen et al., 2012). Their profile in signaling error in expectation of positive versus aversive stimulus associations is less clear. To investigate this, after demonstrating that VTA GABA neurons actively encode conditioned and unconditioned events of either valence, we measured their response to omissions of expected reward or shock. Rats were trained on reward conditioning as described above, where a CS+ predicted the delivery of Ensure solution. In a test session, this reward was omitted on 50% of CS+ trials, intermixed with normal trials (Fig. 3A). As above, we found strong reward-evoked GABA signals. Reward omission resulted in a depression in GABA neuron activity relative to the reward signal that was time-locked to the period of expected reward (Fig. 3C,D; AUC paired t test: t(7)=5.216, p=.0012; min/max z-score in 10-s window after omission: t(7)=9.60, p<.0001). This omission signal dipped significantly below zero (one-sample t test vs 0, t(7)=6.99, p=0002).

**Fig 3.**
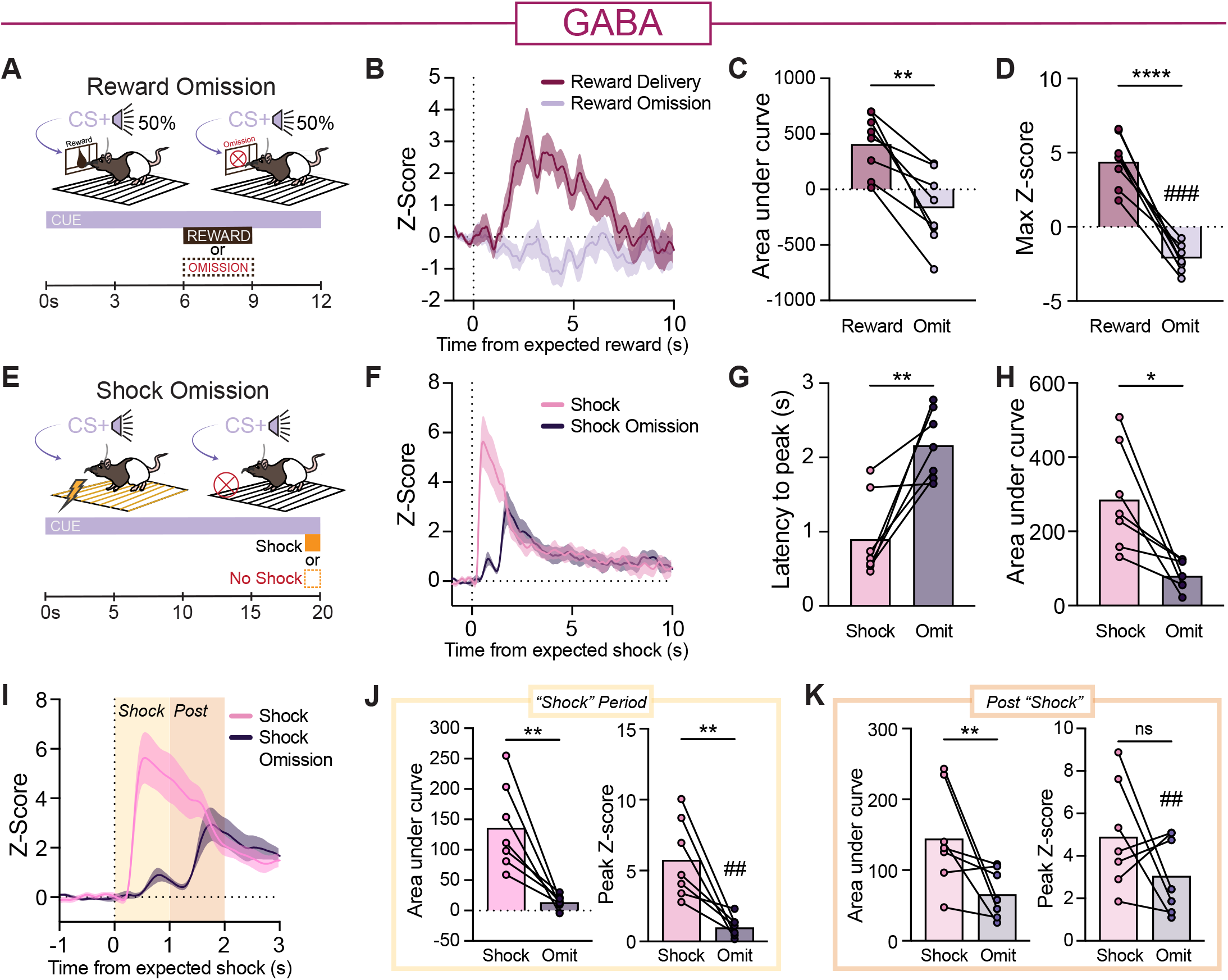
VTA GABA neurons differentially track appetitive and aversive expectation violations. **A)** Schematic of reward omission test. Reward was omitted following 50% of CS+ presentations. **B)** VTA GABA neurons (n=8 rats) showed robust response to reward consumption and a decrease in activity during omission of expected reward. **C)** AUC and **D)** peak measures showed a significant decrease in activity in the omission condition, relative to the reward condition. **E)** Schematic of shock omission test. **F)** VTA GABA neurons (n=7 rats) showed increased activity to shock omission, but with a different pattern than to shock itself. This resulted in a significantly **G)** longer latency to peak signal and a **H)** smaller AUC in the shock omission condition relative to shock. **I)** Zoom in on the trace shown in **F**, highlighting the two-peak shape of the GABA neuron response in the shock omission condition, relative to shock. **J)** During the expected shock window, the GABA neuron response was smaller in the omission condition, relative to the shock condition, for AUC and peak measures. **K)** In the post shock expectation window, the omission signal was smaller for AUC, but not peak measures, relative to the shock condition. Data reflect subject means +/- SEM or individual subjects data. *p< 0.05, **p≤ 0.01, ***p≤ 0.001, ##p<.01 relative to 0, ###p<.001 relative to 0.

Rats were next trained in fear conditioning, as above, where a CS+ predicted the delivery of foot shock. In a subsequent test session, shock was omitted following CS+ presentations (Fig. 3E). Similar to our previous data, footshock evoked robust VTA GABA neuron activity, characterized by a large signal peak during the shock delivery. Omission of this expected shock produced a different pattern of activity, characterized by a small positive peak during the expected shock window and a larger second peak in the period immediately after the expected shock (Fig 3F). Comparing the shock and shock omission conditions, we saw that the latency to the largest GABA signal was significantly longer for omissions (Fig. 3G, t(6)=4.25, p=.0054), and the omission response was smaller overall (Fig. 3H; t(6)=3.42, p=.014). We then directly compared the size of GABA signals in the shock and shock omission conditions, focusing on two time windows reflecting the 1-sec period when shock was expected and the subsequent 1-sec period after expected shock (Fig. 3I). For the expected shock period, GABA signals in the omission condition were smaller for AUC (Fig. 3J; t(6)=4.413, p=.0045) and peak measures (Fig. 3J; t(6)=4.062, p=.0066). For the post shock window, the omission signal was smaller than the shock signal for AUC (Fig. 3K; t(6)=2.485, p=0475), but was not different for the peak measure (Fig. 3K; t(6)=1.37, p=.219). Notably, signals during the shock omission (Fig 3I, purple trace), comprised of two distinct peaks, each were significant above zero (peak 1: one sample t test vs 0, t(6)=3.878, p=.0082; peak 2: t(6)=4.52, p=.004), and the second peak was significantly larger than the first (paired t test peak 1 vs 2: t(6)=2.774, p=.032).

These experiments demonstrate that at the population level, VTA GABA neurons dynamically encode valenced expectations and signal errors when those expectations are violated. These patterns of activity underscore the notion that GABA neurons are engaged in parallel, but uniquely to dopamine neurons during learning.

### VTA GABA but not dopamine neurons integrate valence to encode reward seeking in the face of scaling cost

Above, we show that dopamine and GABA neurons respond similarly in rewarding contexts, but that in aversive situations GABA neurons are uniquely recruited. In each of these previous tasks, the acute learning context was monovalent (appetitive or aversive), and that remained consistent across training. Critically, decision making in natural environments is often within a multivalent context, composed of changing rewards and costs that must be weighed and integrated to guide behavior. Assessing the role of VTA activity in monovalent contexts, as with many previous studies, likely overlooks some of the signaling complexity that must occur when positive and negative variables are simultaneously present and unique computations are required to enable effective decisions. To explore this, we recorded dopamine and GABA neurons during a new motivational conflict task, adapted from our previous studies (Saunders et al., 2013). In this paradigm, rats were required to weigh ongoing, escalating costs when responding to cues and deciding to engage in reward-seeking behavior. Rats were first trained on the reward conditioning paradigm described above. Next, we introduced an ‘aversive barrier’ by constantly electrifying the floor bars directly in front of the reward port, while the back of the chamber remained shock free (Fig. 4A). The intensity of this constant shock escalated across 8 sessions (range: 0.1 mA to 0.4 mA), requiring rats to reconcile competing motivations of seeking reward and avoiding footshock. As this antecedent footshock cost increased across sessions rats overall decreased their cue-driven reward seeking (Fig. 4B; two-way ANOVA, session main effect, F(2.667, 90.69)=61.03, p<0.0001; interaction F(7,238)=37.35, p<0.0001). While rats continued overall to make select port entries and consume rewards, a smaller proportion of port entries were made specifically during the CS+ period (Fig. 4C; paired t test, t(14)=2.54, p=.024), and rats on average stayed farther away from the reward port under higher cost conditions (Fig. 4D).

**Fig 4.**
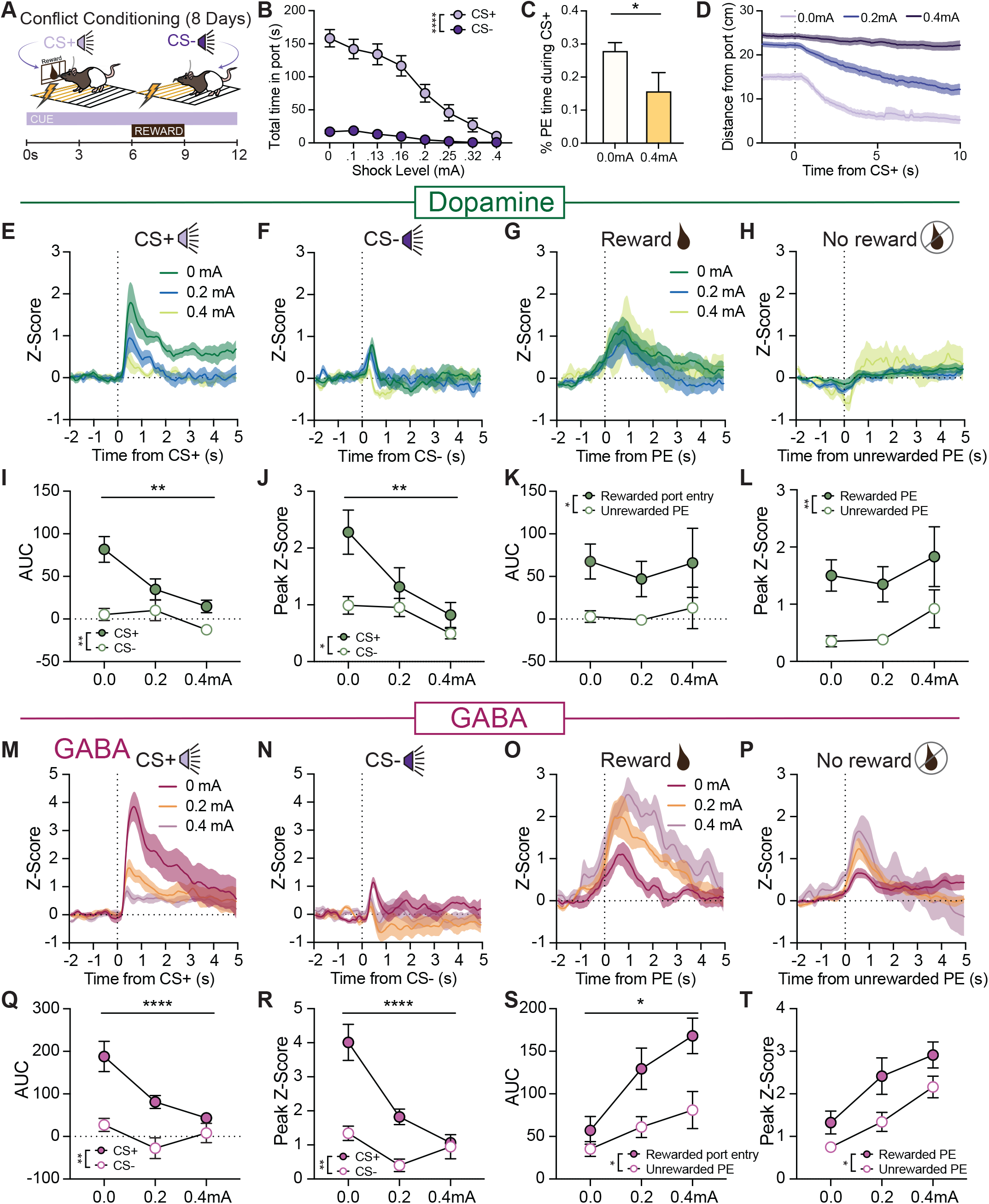
VTA GABA but not dopamine neurons integrate valence during reward seeking in the face of scaling cost. **A)** Schematic of experimental design. Rats (N=18) received cue (CS+) and reward (Ensure) pairings and a separate neutral cue (CS-). After an initial baseline session (0 mA), a cost to reward seeking was introduced in the form of an aversive “barrier”. The floor bars in front of the reward port were continuously electrified and shock intensity was gradually escalated across successive sessions. Reward consumption thus required experiencing footshock. **B)** This resulted in a reduction in reward seeking during CS+ presentations **C)** The proportion of time spent in the reward port during the CS+ decreased with cost, although rats continued to make select port entries and consume rewards. **D)** Analysis of approach behavior showed that rats were less likely to rapidly enter the shock zone in response to the cue at higher shock levels. **E-L)** Calcium recordings from VTA dopamine neurons (n=11). **E)** Dopamine neurons had large CS+ evoked responses initially which reduced as cost increased, **F)** while showing minimal, consistent responses to the CS- cue across cost level. **G)** Reward consumption evoked consistent dopamine neuron activity across cost levels, **H)** while unrewarded port entries (PEs) were associated with minimal dopamine activity. **I)** Dopamine neuron discrimination of CS+/CS- cues decreased as cost escalated, as measured by AUC and **J)** peak measures of the GCaMP signal. **K)** Dopamine neuron activity discriminated rewarded versus unrewarded port entries but there was no change in the pattern of signal as cost escalated for AUC or **L)** peak signal measures. **M-T)** Calcium recordings from VTA GABA neurons (n=7). **M)** GABA neurons had large CS+ evoked responses initially which reduced as cost increased, **N)** while showing small, consistent responses to the CS- cue across cost level. **O)** GABA neuron activity evoked by reward consumption increased as cost escalated, as did **P)** activity in response to unrewarded port entries. **Q)** Average GABA neuron discrimination of CS+/CS- cues decreased as cost escalated, as measured by AUC and **R)** peak measures of the GCaMP signal. **S)** GABA neuron activity discriminated rewarded versus unrewarded PEs, but these signals increased as cost escalated for AUC and **T)** peak signal measures. Data reflect subject means +/- SEM. *p< 0.05, **p≤ 0.01, ****p≤ 0.0001.

Dopamineneuronactivity(Fig.4E-L)tothe CS+weakened as the cue itself drove less reward-seeking behavior in response to growing cost, as measured by AUC (Fig. 4I; cost phase by cue interaction, F(1.86,18.58)=6.71, p=.0073; main effect of phase, F(1.66,16.64)=10.75, p=.0017) and peak signal (Fig. 4J; phase by cue interaction, F(1.61,16.1)=8.345, p=.0049; main effect of phase, F(1.55,15.5)=16.11, p=.0003). Dopamine activity was higher during rewarded port entries compared to unrewarded port entries, however, the pattern of activity did not change in response to seeking under increasing shock intensity for AUC (Fig. 4K; phase by response type interaction, F(1.1,8.27)=.130, p=.551; effect of response type, F(1,10)=6.55, p=.028) or peak measures (Fig. 4L; phase by response type interaction, F(2,15)=.124, p=.88; effect of response type, F(1,10)=11.17, p=.008).

GABA neuron activity (Fig. 4M-T) also decreased on average to the CS+ as the cost escalated and reward seeking became less tied to the cue for both AUC (Fig. 4Q,R; phase by cue interaction, F(1.73,10.38)=34.01, p<.0001; effect of phase, F(1.4,8.38)=9.202, p=.01) and peak (Fig. 3S,T; phase by cue interaction, F(1.86,11.17)=30.97, p<.0001; main effect of phase, F(1.48,8.9)=14.82, p=.0023). In contrast to dopamine neurons, however, we found that the GABA reward signal escalated with rising costs. GABA activity discriminated the rewarded from unrewarded port entries, and this signal scaled at higher levels of shock, compared to sessions of medium and no shock for AUC (Fig. 4S; phase by response type interaction, F(1.13,4.51)=9.48, p=.03; effect of phase, F(1.37,8.24)=16.06, p=.0024) and peak (Fig. 4T; phase by response type interaction, F(1.38,5.52)=2.195, p=.198; effect of phase, F(1.5,9.06)=22.22, p=.0005). In control studies, we determined that these GABA signals do not simply represent changing perceptual salience, as different intensities of unexpected reward (AUC, paired t test, t(6)=1.18, p=.281) or shock (AUC, paired t test, t(6)=0.65, p=.542) did not evoke different levels of GABA neuron activity (Supplemental Fig. 3). This indicates that GABA scaling in our conflict task likely does not reflect a simple readout of stimulus intensity, but rather is an index of motivational state.

### VTA GABA neuron activity encodes shifts in motivational state under conflict

An important feature of our conflict task is that it allows for assessment of the motivated decision state that precedes seeking in the face of a known cost that must be surmounted before receipt of reward. This is distinct from paradigms where a negative stimulus is presented as an unavoidable outcome or punishment after a cue or reward seeking action, or active avoidance paradigms, which likely all reflect somewhat distinct psychological processes (Bornhoft et al., 2025; Bravo-Rivera et al., 2014; Jacobs & Moghaddam, 2020; Lefner & Moghaddam, 2025; Oleson et al., 2012).

To better understand neural encoding of this conflicted decision state, we separately analyzed CS+-evoked calcium signals on trials where rats either engaged in a reward seeking response (approach and port entry) during the cue, or did not (Fig. 5A). We found that cue-evoked GABA, but not dopamine neuron activity, distinguished this cue-evoked conflict state. Dopamine neuron CS+ signals were similar for trials with or without a port entry (Fig. 5B,C; paired t-test; peak t(8)=1.82, p=.106). In contrast, the GABA neuron response to the CS+ was greater on trials when rats subsequently entered the shock zone to make a reward seeking response (Fig. 5D,E; paired t-tests; peak t(5)=4.34, p=.0074). This suggests that GABA neuron responses in this paradigm are not simply a readout of the aversive state/shock experience, but are recruited under situations of decision conflict, to prime reward seeking.

**Fig 5.**
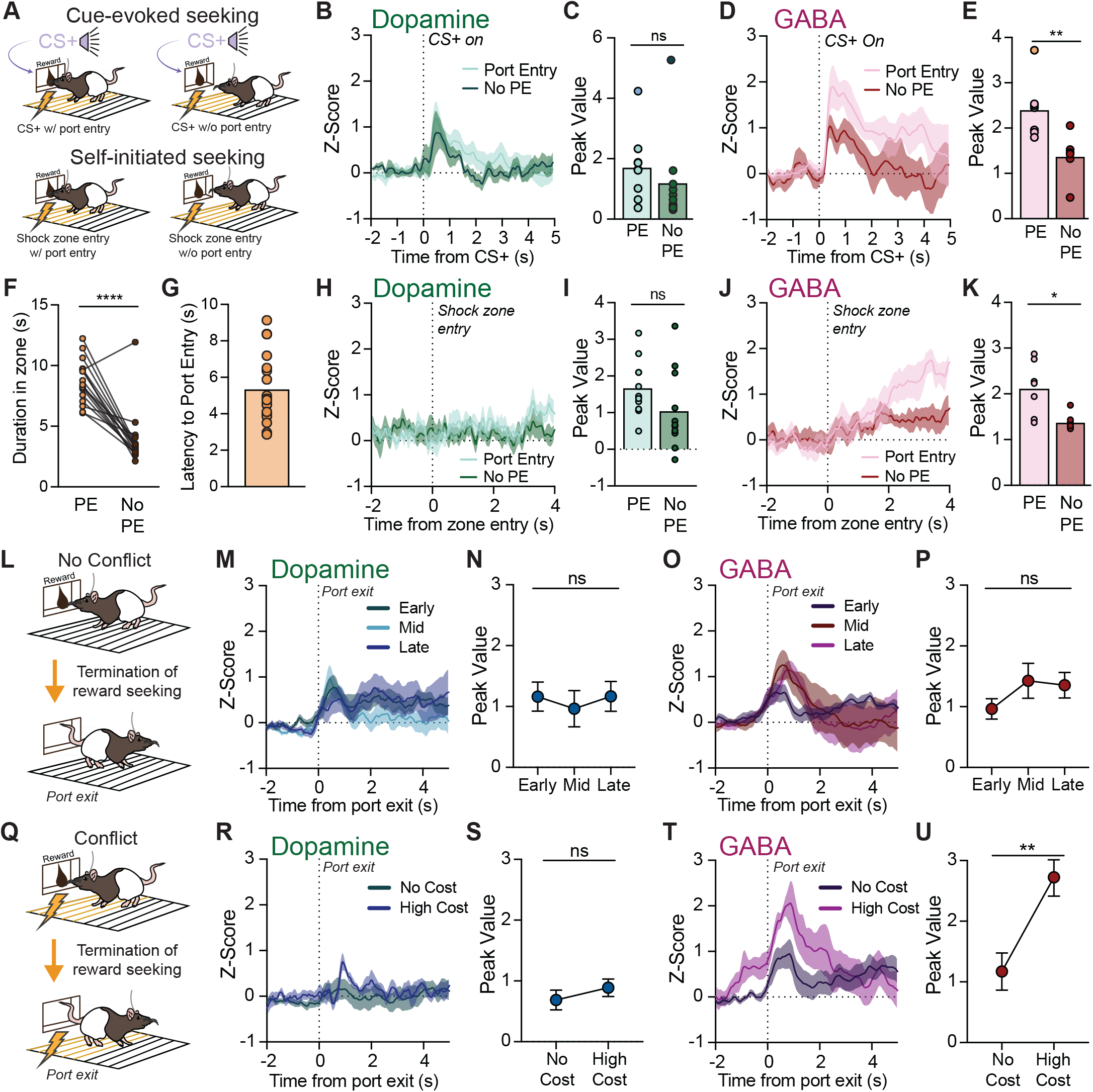
VTA GABA neuron activity encodes shifts in motivational state under conflict. **A)** In the conflict task (N=18), we examined reward seeking events (approach and entry to the reward port) from the perspective of cue onset (cue-evoked), or self-initiated movements. **B-E)** Dopamine and GABA neuron cue signals were separately analyzed on trials where a port entry response did or did not occur, focusing on the 0.2 mA cost session. **B**,**C)** CS+-evoked dopamine neuron activity did not discriminate reward seeking under cost. **D**,**E)** In contrast, GABA neuron activity evoked by the CS+ was higher on trials where a PE was subsequently made. **F-K)** Self-initiated visits into the shock zone in front of the reward port were calculated from behavioral video tracking. Dopamine and GABA neuron activity was compared for zone visits that did or did not include a port entry. **F)** Shock zone visits were longer in duration when a port entry was made, compared to visits that did not include a port entry. **G)** The average latency between shock zone entry and port entry was around 5 seconds. **H**,**I)** Dopamine neuron activity was similar for shock zone entries that did or did not lead to a port entry. **J**,**K)** GABA neuron activity was higher following shock zone entries that lead to a port entry, compared to zone entries without a port entry. **L-U)** Dopamine and GABA neuron signals were time locked to the termination of port entries (port exit) for **L)** reward conditioning and **Q)** conflict conditioning. **M**,**N)** Dopamine neurons and **O**,**P)** GABA neurons showed a small port entry exit signal that was stable across reward conditioning. **Q)** Port exit signals in conflict conditioning were compared for a no cost session and a high cost session. **R**,**S)** Dopamine neurons showed minimal response to port exit, regardless of the presence of cost. **T**,**U)** GABA neurons, in contrast, showed scaled activity associated with port exit in the high cost condition. Data reflect subject means or individuals +/- SEM. *p< 0.05, **p≤ 0.01, ****p≤ 0.0001.

Beyond this cue-driven motivation, we also saw an increase in self-initiated port entries. This decoupling of behavior and cue was not unexpected, as the cue itself was less of a driver of reward seeking in situations with increased cost. To understand the pattern of neural activity in relation to self-initiated seeking events, we incorporated DeepLabCut-based pose estimation analysis to identify the periods when rats made entries into the shock zone. While there were overall consistent patterns of shock zone behavior across rats as cost increased, individual rats had variable responses to different shock levels, possibly reflecting different perceptions of a given cost level. To get a better sense of the periods of highest decision conflict preceding reward seeking, for each rat, we compared the conflict session that produced behavior levels that were 10% of that animal’s behavior on the zero shock/no conflict session. Thus, we isolated the period when rats volitionally made the decision to enter the shock zone and make a port entry, compared to other entries that are more exploratory (Fig. 5A). At this high “perceived cost”, shock zone visits were longer in duration when rats made a port entry, compared to when they exited the zone without entering the port (Fig 5F: paired t test, t(16)=7.285, p<.0001). Following zone entries, the latency to make a port entry, when they did occur, was around 5 seconds (Fig. 5G). Using this as a threshold, we looked at dopamine and GABA neuron signals in the first few seconds following the initiation of movements leading to shock zone visits, comparing visits that led to port entries versus visits that did not produce a port entry. Dopamine neuron activity did not differ on these two types of shock zone visits (Fig. 5H,I: signal peak, t(10)=2.036, p=.0691). In contrast, GABA neuron activity was significantly greater after movement initiation in cases where shock zone entry was accompanied by a reward seeking response (Fig. 5J,K: signal peak, t(6)=3.598, p=.0114), a pattern reflective of the intention to seek reward in the face of high perceived cost. This adds further support to the conclusion that GABA neuron signals encode reward seeking under conflict, and do not simply read out shock level/aversion. Notably, this rise in GABA signal was temporally distinct from the CS+ and reward port entry scaling we show above, indicating that GABA neuron activity encodes multiple components of reward seeking under conflict.

We next examined the neural activity profiles associated with the termination of reward seeking events, comparing reward conditioning with and without associated cost. Calcium traces were time-locked to the end of port entries made during reward conditioning, across early (day 1), middle (day 8 or 9), and late (day 13 or 15) training phases (Fig. 5L). Dopamine neurons showed a small bump in activity at port exit, which was consistent in size across training phases (Fig. 5M,N: one-way ANOVA, no effect of training phase, F(1.8,17.2)=.2604, p=.753). GABA neurons showed a similar profile, with consistent port exit signals across reward conditioning (Fig. 5O,P: no effect of training phase, F(1.9,13)=1.42, p=.275). We applied the same analysis to port exits made during conflict conditioning, comparing the 0mA no cost session to the highest cost session reached by each rat (Fig. 5Q). Dopamine neurons again showed a minor bump in activity at the termination of reward seeking, which did not change when cost was present (Fig. 5R,S: paired t test, t(10)=1.09, p=.311). GABA neuron activity, however, was higher at port exit when cost was present (Fig. 5T,U: effect of cost, t(6)=4.03, p=.0069). Thus, VTA GABA neuron activity, but not dopamine neuron activity, tracks the initiation and termination of reward seeking events preferentially under conditions of decision conflict.

Collectively, the results from this set of studies indicate that VTA GABA neurons perform a unique computation during reward seeking, compared to dopamine neurons, encoding externally or internally generated shifts in motivational state as the ongoing cost to seeking rises, and to enhance the representation of rewards that are experienced under threat.

Cue-evoked VTA dopamine and GABA neuron activity differentially predicts movement vigor across appetitive and aversive learning contexts

As a final look at differences in behavioral encoding for dopamine and GABA neurons, we next explored the connection between cue-evoked activity in each population and movement invigoration late in conditioning across each learning paradigm (Supplemental Figure 5). Making use of our pose estimation pipeline, we calculated rat’s speed during cue presentations. Performing correlations between the area under the curve of CS+ cue-evoked signal and these variables, we found that cue-evoked dopamine neuron activity was predictive of movement vigor on a trial-by-trial basis, only in reward conditioning. Dopamine signals were negatively correlated with the latency to reach maximum speed after cue presentation (Suppl Fig 5A: Pearson correlation, p<.0001), and positively correlated with the average speed reached during the reward cue (p=.014). Dopamine neuron activity was not correlated with speed for fear conditioning (Suppl. Fig. 5B: p=.1908) or conflict conditioning (Suppl. Fig. 5C; p=.1824). In contrast, for GABA neurons, there was no correlation between cue-evoked activity and speed during reward conditioning (Suppl. Fig. 5D: latency to max, p=.4674; average speed, p=.4901), but we saw trends for correlations between GABA neuron activity and speed in fear (Suppl. Fig. 5E: p=.0865) and conflict conditioning (Suppl. Fig. 5F: p=.0722).

This analysis indicates that GABA neuron activity is less connected to conditioned vigor than dopamine in reward learning contexts, but vigor-related signals in GABA neurons may come online when aversive stimuli are present.

### Inhibition of VTA GABA neurons disrupts motivation to seek reward in the face of cost

Given the specific recruitment of VTA GABA neurons during conflicted reward seeking, we predicted that inhibiting this population would decrease reward seeking under cost. To test this, we expressed the inhibitory actuator halorhodopsin in VTA GABA neurons, for optogenetic manipulation (Fig. 6A-C). Control rats received a virus coding for YFP only. Rats were trained on a modified version of the motivational conflict task. In the acquisition phase (7 sessions), they learned to make active nose pokes to self-administer sucrose reward (Fig. 6D,E; main effect of nose poke, F(1,11)=46.40, p<.0001). On a subset of subsequent sessions, a constant electric barrier “cost” was introduced directly in front of the nose pokes and reward port (Fig. 6F). Sessions with cost were then intermixed with sessions with no cost present. Rats made significantly fewer active nosepokes during sessions with cost compared to no cost sessions (Fig. 6G; paired t test, t(11)=6.29, p<.0001).

**Fig 6.**
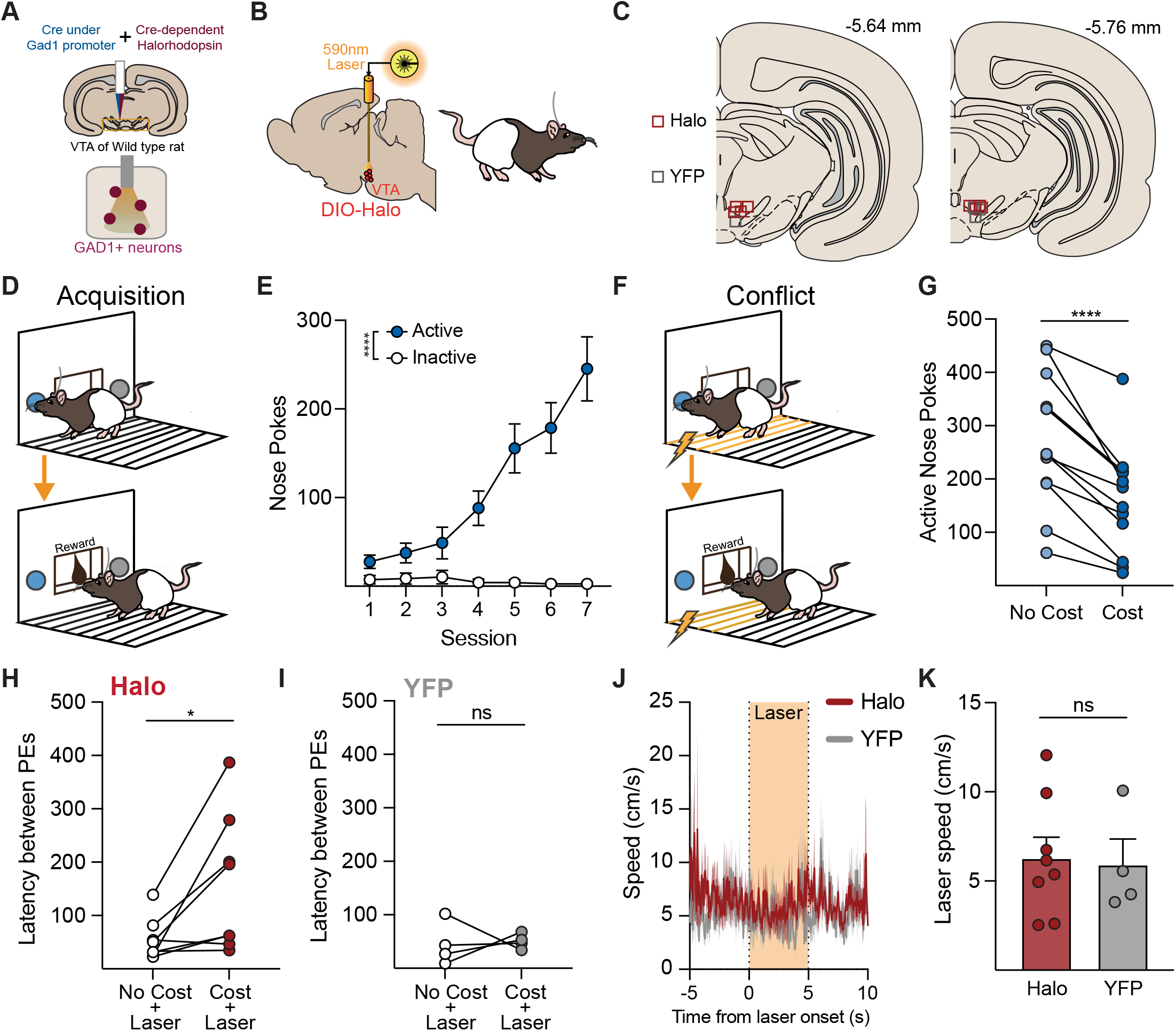
VTA GABA neuron inhibition disrupts motivation to seek reward in the face of cost. **A)** Approach for targeting halorhodopsin to VTA GABA neurons. **B**,**C)** Optic fibers were targeted to the VTA for delivery of orange (590-nm) for optogenetic manipulation. **D)** Acquisition: Rats (N=12) were trained to nose poke to self administer sucrose reward, **E)** reliably discriminating active from inactive nose pokes responses. **F)** Conflict: A cost to reward seeking was introduced via electrification of the floor bars directly in front of active nose poke and reward port. **G)** Rats made fewer active nose pokes on sessions where cost was present, compared to interleaved no cost sessions. On test sessions where cost was or was not present, laser (5-s constant pulse, 4 times per session) was delivered to the VTA, programmed to occur in between ongoing reward seeking bouts. **H)** For rats expressing Halo (n=8), optogenetic inhibition of VTA GABA neurons significantly increased the latency between reward seeking responses when laser was delivered in the presence of the electric barrier cost, relative to no cost laser sessions. **I)** For YFP control rats (n=4), laser delivery had no effect on reward seeking. **J)** Laser delivery had no effect on acute movement speed, and **K)** Halo and YFP rats did not differ for average movement speed. Data reflect subject means or individuals +/- SEM. *p< 0.05, ****p≤ 0.0001.

Previous work suggests that optogenetic inhibition of VTA GABA neurons following an action can mildly reinforce behavior, particularly if inhibition lasts for longer than a few seconds (Corre et al., 2018). Our photometry data above indicates that VTA GABA neurons are active at various timepoints preceding reward seeking under cost, suggesting that this signal is important for marshalling motivation to seek amid conflict. Based on this information, we chose a conservative approach for optogenetic inhibition. Laser was delivered in brief 5-sec windows at four time points spaced throughout each test session, programmed to not overlap with an ongoing seeking action. Laser was delivered in this fashion on test sessions in the presence of the electric barrier cost and in comparison sessions where the cost was not present. To look at the impact of this manipulation, we calculated latency between reward port entries in the periods following laser delivery. For the active virus (Halo rats), optogenetic inhibition of VTA GABA neurons resulted in a significantly greater latency between port entries specifically when laser was delivered on sessions with cost present, compared to laser sessions without cost (Fig. 6H; paired t test cost vs no cost; t(7)=2.80, p=.0267). In contrast, we saw no effect on reward seeking for the control YFP group (Fig. 6I; paired t test cost vs no cost, t(3)=.197, p=.856). We next examined general locomotion during the laser delivery windows, as previous studies have reported that optogenetic manipulation of VTA GABA neurons can influence movement (van Zessen et al., 2012). Here, we found no impact of laser delivery on acute movement (Fig 6J), and Halo and YFP groups showed the same average speed during laser periods (Fig 6K; unpaired t test, t(7)=.196, p=.85). Together with our recording data above, these studies indicate that VTA GABA neuron activity encodes reward seeking motivation and is functionally recruited to drive reward seeking under motivational conflict.

## Discussion

Our studies reveal that GABA neurons actively encode appetitive and aversive cues and outcomes and, critically, integrate the experience of both valences to guide reward seeking under motivational conflict. Dopamine neurons had a narrower encoding profile in contrast, consistently responding only to appetitive stimuli. Our results provide strong support for evolving notions of VTA functional heterogeneity, suggesting that GABA neurons contribution to appetitive motivation is not necessarily in opposition, but is unique and in parallel with dopamine neuron signaling, serving to consolidate valence calculations to promote motivation in multivalent decision-making states. Compared to dopamine neurons, our results indicate that GABA neurons reflect a broader learning signal that is important for directing appropriate behavioral responses in complex environments. Together, our data offer insight into a number of ongoing questions regarding the role of midbrain systems in controlling different facets of learning, as well as the heterogeneous and dynamic nature of the VTA.

We found that VTA GABA neuron activity encoded cost in multiple ways: by scaling reward outcome encoding with escalating cost, and through increased activity preceding when rats made reward seeking actions in the face of cost. Further, we found that GABA neuron activity is necessary for normal reward seeking when decision conflict is present. Collectively, this supports an important new insight into GABA neuron function, indicating that they do not merely represent aversive experiences, but are critical for marshaling reward-seeking motivation and signaling the outcome of actions made under decision conflict. This positions VTA GABA neurons with an important motivational function that is independent of or at least in parallel to dopamine neurons. Interestingly, we find that dopamine neurons were relatively insensitive to threat or cost, with minimal changes in negative outcome encoding, and no clear connection between cue-evoked signals and the choice to encounter cost. A relative lack of aversion-mediated signals in our data is on its own and important conclusion. Our data emphasize that at the population level, dopamine neurons are biased for responsiveness in appetitive contexts, while there is heterogeneity among specific subpopulations that remains to be fully understood (Lammel et al., 2014; Lopez & Lerner, 2025; Ungless et al., 2010).

Notably, GABA neurons discriminated between the outcome-predictive cue and the neutral cue most clearly in the appetitive context but not the aversive context. This suggests that VTA GABA neurons may have a prepotent bias towards threat encoding, which could prime them to integrate negative valence into reward computations. This is consistent with our conflict paradigm recordings, where we saw elevated GABA neuron activity when unrewarded port entries were made under cost, and in response to the termination of reward consumption events under cost. Further, our optogenetic inhibition study shows that VTA GABA neuron activity is especially important for normal reward seeking when animals are faced with a conflicted motivational state. Collectively this supports a framework where VTA GABA neurons are more reactive to new information - in this case, the presence or absence of reward - when threat or conflict is in play. This conclusion helps consolidate past findings that show evidence for both an “anti-reward” function and an essential reward prediction learning calculation in VTA GABA neurons (Eshel et al., 2015; Tan et al., 2012).

Our studies demonstrate that VTA GABA neurons can signal errors when expectations are violated, with different patterns for appetitive and aversive associations. This is distinct from some past research reporting minimal change in GABA neuron activity during reward omission, in single-unit recordings in head fixed mice (Cohen et al., 2012; Eshel et al., 2015). We also did not see strong evidence of GABA neuron ramping activity preceding reward delivery in the simple reward conditioning task, like in those past studies. There are multiple possibilities explaining these differences. It is likely that midbrain neuron encoding properties differ for freely-moving animals, in our case here, given there is an increase in the variability of specific timing that animals interface with rewards. Our results also leverage fiber photometry methods, which allow for assessment of bulk activity measurements. While it is likely that some individual GABA neurons exhibit ramping activity before expected reward, our results suggest that it might not be the average output of this population. In our conflict paradigm, we did see an increase in GABA neuron activity as rats approached the reward port, but only when cost was present. Thus, expectation-related signals in VTA GABA neurons may be more readily engaged under learning conditions when risk, cost, or uncertainty are at play. That we see opposing error signals for reward and shock omission suggests that VTA GABA neurons track different components of value and motivational salience. Taken together with our conflict recordings, these results underscore the integrative nature of the VTA GABA signal. Given that individual dopamine neurons are inconsistently engaged by aversive stimuli, it has been difficult to incorporate aversive learning states into comprehensive frameworks of dopamine function (Bromberg-Martin et al., 2010; Gershman et al., 2024; Lee et al., 2024; Schultz, 2013). Our studies motivate including local GABA neurons more prominently into the parameter space of computational models of the VTA. In particular, our data suggest that GABA neurons operate not just as regulators of dopamine neurons, but also as parallel processors by uniquely incorporating cost or risk into calculations of motivation and value.

In contrast to previous findings showing that VTA GABA activation results in acute VTA dopamine inhibition (Tan et al., 2012), but consistent with other data (Elum et al., 2024; Lefner & Moghaddam, 2025; Tolu et al., 2013), our results show that these two populations, recorded from the same area of the VTA, are active at the same time. Further work is needed to understand this from a physiological and circuit level. Ex vivo work indicates that VTA GABA and dopamine neurons do not have a single mode of interaction, but the firing rate, patterning, and synchrony of activity in GABA neurons can have different influences on dopamine neurons. The complexity of VTA microcircuitry is particularly reflected in GABA neurons, which encompass local and distal projections and modulate dopamine through direct inhibition, poly-synaptic disinhibition, and amplification of phasic firing (Bocklisch et al., 2013; Lobb et al., 2010; Morozova et al., 2016; Ostroumov et al., 2016; Stamatakis et al., 2013; Tan et al., 2012; Tolu et al., 2013). This interplay makes it possible that concurrent dopamine and GABA neuron activity, as our data imply here, could uniquely shape the pattern of specific sub-population of dopamine neurons without altering population level dopamine activity. This helps contextualize earlier studies that show a prominent inhibitory effect of VTA GABA neurons on dopamine neurons (Tan et al., 2012), and offers mechanistic space for parallel and collaborative GABA/dopamine activity profiles. One interesting possibility our data suggests is that the level of engagement of dopamine or GABA neurons in the VTA is titrated based on the extent of decision conflict currently required by the environment.

Notably, stress can engage unique plasticity in VTA GABA neurons, altering their firing properties and changing the way they can interface with dopamine neurons (Bouarab et al., 2019, 2025; Lowes et al., 2021; Ostroumov et al., 2016). Given the mild nature of the cost in our task and ability for rats to avoid it, we suspect our effects are unique from many stress manipulations, but this comparison raises the interesting possibility that conflicted motivational states may engage a different “mode” for VTA GABA neurons. This could be a possible explanation for the unique activity profiles we see here under conditions of cost. However, given that we show VTA GABA neurons respond similarly to dopamine neurons in the absence of stress or aversive stimuli during reward learning, aversive experience is not necessary to engage parallel learning-related VTA GABA neuron activity. Our results highlight the complexity of valence encoding in the VTA, and motivate future studies involving an integration of appetitive and aversive valence conflict, which likely engages unique neuronal profiles from many classic studies. Our conflict paradigm, as well as other avoidance learning and valence discriminations tasks (Bornhoft et al., 2025; Bravo-Rivera et al., 2014; Jacobs & Moghaddam, 2020; Lefner & Moghaddam, 2025; Oleson et al., 2012), offer some possible ways to progress on this front.

Our results underscore other emerging questions surrounding VTA heterogeneity. We used common markers to target our populations of interest, but there is rich genetic heterogeneity across VTA dopamine and GABA neurons (Miranda-Barrientos et al., 2021; Morales & Margolis, 2017; Olson & Nestler, 2007; Paul et al., 2019; Poulin et al., 2018; Simon et al., 2024). In our comparison studies, VTA calcium recordings using another GABAergic marker, mDlx, also showed similar appetitive and aversive encoding, indicating that this activity phenotype is not specific to Gad1 neurons. Functional heterogeneity among different GABA populations remains a relatively open area of investigation (Koutlas et al., 2024; Margolis et al., 2012; Root et al., 2020; Wang et al., 2024). VTA GABA neurons have several non-dopamine projection targets outside of the VTA, with unique roles (Al-Hasani et al., 2021; Breton et al., 2019; Oriol et al., 2024; Taylor et al., 2014; Yang et al., 2018; Zhou et al., 2022). The detailed function of VTA GABA projections is just beginning to be explored, and the anatomical and subregional complexity introduces challenges. It remains unclear, for example, if there are distinct VTA GABA ensembles that preferentially respond to positive valence or negative valence, and/or uniquely engage local dopamine neurons. Recent work also suggests that many, if not most individual VTA GABA neurons make both local and distal connections (Oriol et al., 2024), an organization that would set the stage for complex GABA-mediated regulation of reward and aversion systems in the brain. Future studies combining genetic dissection of these populations with single unit imaging or physiology will be critical to unpack this complexity.

Our results indicate that VTA GABA neurons have a more nuanced role in conditioned appetitive motivation than previously appreciated. This population acts in tandem with dopamine neurons to promote behavior when animals are faced with reward-seeking choices under conflict. Given the prominence of multi-valent contexts in natural decision making, the pattern we describe for VTA GABA neurons is likely a common encoding profile, rather than the exception. These data point to the need for a strong experimental focus on VTA heterogeneity in complex learning and decision-making frameworks, as well as compulsive disease models.

## Acknowledgements

We thank all members of the Saunders lab for discussions and feedback on this project.

## METHODS

Subjects: Male and female Long Evans rats were used (N=50; 19F, 31M). Dopamine neurons were targeted in TH-cre transgenic rats, expressing Cre recombinase under the tyrosine hydroxylase (TH) promoter (Engel et al., 2024; Witten et al., 2011). GABA neurons were targeted in wild type rats. All rats weighed 250-500g at the time of surgery and were 3-6 months old at the time of behavior and photometry recordings. Rats were paired or singly housed throughout the duration of the experiment in a vivarium with a 12 hour light / 12 hour dark cycle (lights on at 0700, lights off at 1900) with ad libitum access to food and water. During the reward conditioning and motivational conflict phase animals were food restricted - fed once/day to maintain 90% free feeding body weight. All procedures involving animal subjects were in compliance with Institutional Animal Care and Use Committee (IACUC) approval and in accordance with the National Institutes of Health’s animal care guidelines.

### Viral vectors

To target dopamine neurons, a cre-dependent AAV coding for GCaMP8f was injected into the VTA in TH-cre positive rats (750nL, AAV5-syn-DIO-jG-CaMP8f-WPRE; Addgene). To target GABA neurons for fiber photometry, an AAV delivering Cre under the Gad1 promoter (AAV5-Gad1-cre; University of Minnesota Viral Vector and Cloning Core) was simultaneously infused with the cre-dependent GCaMP8f or GCaMP8m into the VTA of wildtype rats (125-250nL Gad1-Cre, 250-750nL DIO-GCaMP), similar to previous studies (Scott et al., 2023; Wakabayashi et al., 2019). A separate set of wild-type rats received VTA-targeted injections of a GCaMP virus under the GABAergic interneuron promotor mDlx (800nL, AAV5-mDlx-GCaMP6f; Addgene). For optogenetic experiments, a cre-dependent AAV coding for halorhodopsin (500 nL, AAV5-DIO-eNpHR3.0-YFP) paired with the Gad1-Cre virus or an AAV delivering eYFP only was infused into the VTA to selectively target midbrain GABAergic neurons.

### Stereotaxic surgery

Rats were induced under 5% isoflurane anesthesia and maintained at 1-3% for the duration of the surgery. Rats received carprofen (5 mg/kg), cefazolin (70 mg/kg), and saline subcutaneously at the beginning of the surgery. An incision was made to reveal the top of the skull and holes were drilled for viral delivery (−5.6 A-P, -/+0.8 M-L, -8.2 D-V), optic fiber placement (−5.6 A-P, -/+0.8 M-L, -8.0 D-V), and skull screws. All coordinates are in mm relative to bregma and skull surface. Viruses were delivered at a rate of 0.1μL/min. Once infusion finished, the needle was raised 100 μm and left to sit for 10 minutes before being fully removed. Optic fiber implants (9mm length, 400μm diameter, Doric Lenses) were lowered into the VTA and secured in place with dental acrylic around the skull screws. Topical anesthetic and antibiotics were applied to the incision area and the rat was monitored on a heating pad in a sterile home cage until fully alert and awake. Rats were given carprofen and cefazolin for three days following surgery and weighed with health evaluations for 6 days. Rats recovered for at least 4 weeks following viral injection before beginning experiments. Final anatomical placements in the VTA ranged from 0.5mm to 1.1mm lateral from Bregma, with a mean of +.75. We found no correlation between recording position and cue or outcome-related photometry signals, for either neuron type.

### Fiber photometry recordings

To assess Ca2+ dynamics in the VTA during Pavlovian conditioning, we measured GCaMP fluorescence using fiber photometry. For all behavioral experiments, rats were tethered to a low autofluorescence optic cable sheathed in a lightweight armored jacket (Doric-Lenses). Some behavioral sessions did not record photometry signals to avoid photobleaching the GCaMP fluorescent proteins. Recording data included here are sampled from the early (Day 1), middle (Days 8 or 9), and late (Day 13 or 15) of reward conditioning. On photometry recording days, 415 nm and 465 nm LEDs (Doric-Lenses) set at 50μW were delivered via a fluorescence mini-cube (Doric-Lenses). The isosbestic (415) and Ca2+ dependent (465) channels were sinusoidally modulated at 211Hz and 330Hz respectively. Fluorescence from the implanted optic fiber was transmitted through the mini-cube to be filtered, amplified, and focused onto a high sensitivity photoreceiver (Newport, Model 2151). A real-time signal processor (RZ5P, Tucker Davis Technologies) using Synapse software modulated the power of the LED output and recorded photometry signals, sampled at 6.1kHz. Events during the behavioral session (e.g., foot shock and cue presentations), were recorded in the photometry data file with a TTL signal time stamp from the MED-PC behavioral program. Videos of each behavioral session were recorded at 10-20 FPS with corresponding time-stamps of each frame in the photometry file.

### Habituation

Before any behavioral testing, animals were habituated to cable tethering and med-associates chambers with 1-2 habituation sessions of 30 minutes with the chamber lights and fans running.

### Reward conditioning

Animals received between 13 and 15 days of reward conditioning. Each session consisted of 20 conditioned stimulus plus (CS+) and 20 conditioned stimulus minus (CS-) presentations with an average inter-trial interval of 80 seconds. The CS+ and CS- were distinct auditory stimuli counterbalanced across animals. The CS+ was 12 seconds long with the onset of a pump providing Ensure reward delivered between seconds 6 and 9. The CS- was 12 seconds long with no additional events. Reward seeking was measured by duration of time in the reward port during the CS+, CS- or inter-trial interval (ITI).

### Fear conditioning

Animals received 3 days of fear conditioning. Each session consisted of 15 conditioned stimulus shock (CS+) and 15 conditioned stimulus minus (CS-) presentations. The CS+ and CS- were distinct auditory stimuli counterbalanced across animals. The CS+ was 20 seconds long with the onset of a 0.4mA footshock delivered during the last second. The CS- was 20 seconds long with no additional events.

### Expectation violation

After initial reward conditioning, in which animals learned to predict reward paired with the CS+, animals received a session in which half of the CS+ presentations were paired with reward delivery (as expected) and the other half had the reward omitted. Rewarded and unrewarded trials were presented in random order. After initial fear conditioning, in which animals learned to predict shock paired with the CS+, animals received an extinction session in which the shock was omitted following each CS+ presentation.

### Motivational conflict task

Animals received 8 days of the motivational conflict paradigm (Adapted from Saunders et al., 2013). Each session consisted of 20 CS+ and 10 CS- presentations. The cues and the pump were the same as during reward conditioning. On sessions 2-8, the 7 floor bars closest to the reward port were electrified with a mild electric shock. The shock ranged from 0.0mA - 0.4mA and increased across the 8 sessions. Reward seeking was measured by duration of time in the reward port during the CS+, CS- or inter-trial interval (ITI).

### Stimulus scaling – reward

Animals received 30 unexpected rewards equally split between 80% or 20% Ensure diluted with water. 80% and 20% Ensure were delivered on opposite sides of the chamber to the right or left port on the first day and swapped for the second day. Photometry traces were time-locked to the first port entry following the pump and averaged across both sessions.

### Stimulus scaling – shock

Animals received 5 unexpected foot shocks of 0.2mA and 0.4mA for 1 second each. The order of shock intensities was counterbalanced across animals.

### Operant sucrose self-administration

Animals underwent 7 60 minute training sessions to learn to self-administer sucrose. Entries into the active port were paired with lights inside the port, white noise, and pump delivery of sucrose for 3 seconds. Entries into the inactive port had no paired outcomes. The floor bars directly in front of the ports and reward port were then electrified to deliver a mild footshock upon entry into the electric barrier zone. The intensity of the shock was modulated across sessions delivering 0.0, 0.07, 0.10, and 0.13, and 0.16mA with this shock “cost” days intermixed with no cost days.

### Optogenetics

590nm light was delivered 4 times per session at 5, 20, 35, and 50 minute timepoints. Light traveled from DPSS lasers, through fiber optic cables, to an intracranial fiber implant leading to the VTA. Laser was delivered at a constant pulse, for 5 seconds at each delivery window. Power was set to 10mW output at the beginning of each session.

### Fiber Photometry Analysis

Recordings were analyzed with a custom MATLAB (Mathworks) pipeline. First, signals were low-pass filtered, downsampled to 40Hz, and a least squares linear fit was applied to the isosbestic channel (415 nm) to align it to the calcium-dependent, 465 nm, channel. The fitted isosbestic channel was then used to normalize the 465 signal with the calculation ΔF/F = (465-nm signal – fitted 415-nm signal)/(fitted 415-nm signal). Individual trial traces were z-scored to the 5s immediately preceding cue, port entry, and footshock events in order to avoid effects of drift across the session. Area under the curve (AUC) values took the z-scored trace and calculated numerical integration via the trapezoidal method using the trapz function (MATLAB). AUC and peak Z-score values were calculated between event onset and 2 seconds immediately after.

### Automated pose-estimation

Markerless tracking of animal body parts was conducted using version 2.2.1.1 of the DeepLabCut (DLC) Toolbox (Mathis et al., 2018) and analysis of movement features based on these tracked coordinates was conducted in MATLAB. DeepLabCut 2.2.1.1 was installed in an Anaconda environment with Python 3.8.4, CUDA 11.7 and Tensorflow 2.10. DeepLabCut Model: 2090 frames from 35 videos (32 different animals, 3 experiments) were labeled and 807 outlier frames were re-labelled to refine the network. Labeled frames were split into a training set (95% of frames) and a test set (5% of frames). A ResNet-50 based neural network (Insafutdinov et al., 2016) was used for 1,030,000 training iterations. After the final refinement we found the test error was 4.1 pixels, the training error was 3.13 pixels and with a p-cutoff of 0.85 training error was 2.99 pixels and test error was 3.68 pixels. The body parts labeled included the nose, eyes, ears, center of head or fiber optic implant, shoulders, tail base, and an additional three points along the spine. Features of the environment were also labeled, including the 4 corners of the apparatus floor, two nose ports, two cue lights, two reward ports, and 3 LED indicator lights when active. DLC coordinates and confidence values for each bodypart and frame were imported to Matlab and filtered to exclude body parts/features from any frame where the confidence was < 0.7. For labeled features of the environment, which have a fixed location, the average coordinates for that recording were used for analysis. To convert pixel distances to the real chamber dimensions, for each video, a pixel to cm conversion rate was determined. The distance (in pixels) between each edge of the environment floor and the diagonal measurements from corner to corner was measured, and these values were divided by the actual distance in cm. The mean of these values was then used as the conversion factor. Movement speed was calculated from the implant coordinates frame by frame using the formula: [distance moved (pix per cm) * framerate] to give movement speed in cm/s.

Freezing was detected by assessing locomotion of the nose, implant, left and right ears, and 4 points spanning the length of the animal’s back. The movement threshold for detecting locomotion was calibrated to animal size using a scale factor determined from the relationship between body size (distance between the shoulder and bottom back point) and the optimal threshold for detecting movement in a separate group of animals not used for this study (n = 4). This threshold was used to detect movements in the face and head, and for remaining body parts this value was multiplied by two to accommodate detection of the finer movements in the face/ head vs larger movements in the body. Freezing bouts were detected when all of the visible body parts were below the respective movement threshold and a sliding window (0.3s) was used to determine when the speed of 2 or more body parts exceeded the movement threshold for the window duration, indicating the beginning and end of a freezing bout. Freezing periods shorter than 1s in duration were excluded and frames in which less than 3 body parts were visible were ignored. The starts and ends of bouts were removed if they consisted only of frames where an insufficient number of body parts were visible. We quantified freezing as the percent of trials with freezing initiated in the last 5 seconds of the CS+ or CS-prior to the footshock or lack thereof (seconds 14-19 post-cue onset).

### Zone analysis

The shock zone was established 10 cm out from the left side of the chamber floor using the conversion factor to scale zone location accurately for each session and animal. Entries began when the mid-point between the shoulder and top back first entered the identified zone for 0.2s, and terminated before this bodypart left the zone for 0.2s. If an ongoing locomotion bout coincided with zone entry, zone entry was realigned to the start of this locomotion bout instead of zone entry.

### Histology

After experiments, rats received i.p. injections of Fatal-Plus (2 ml/kg; Patterson Veterinary) to induce a deep anesthesia, and were transcardially perfused with cold phosphate buffered saline (PBS) followed by 4% paraformaldehyde (PFA). Brains were removed and post-fixed in 4% PFA for ∼24 h, then cryoprotected in a 25% sucrose in PBS for 48 h or until sectioning. The tissue was sliced at 40 microns on a cryostat (Leica CM1900). To confirm viral expression and optic fiber placements, brain sections containing the midbrain were mounted on microscope slides and coverslipped using Vectashield mounting medium containing DAPI counterstain. Fiber tissue damage was then visualized on a Keyence BZ-X710 microscope. Rats were included in analysis only if fiber damage was no more than 500 microns dorsal to the target regions.

### Immunohistochemistry

Sections were washed in PBS and incubated with bovine serum albumin (BSA) and Triton X-100 (each 0.2%) for 20 min. 10% normal donkey serum (NDS) was added for a 30-min incubation, before primary antibody incubation (rabbit anti-TH, 1:500, Fisher Scientific and/or mouse anti-GFP, 1:500, Abcam) overnight at 4 °C in PBS with BSA and Triton X-100 (each 0.2%). Sections were then washed and incubated with 2% NDS in PBS for 10 min and secondary antibody was added (1:200 Alexa Fluor 594 donkey anti-rabbit and/ or 1:200 Alexa Fluor 488 donkey anti-mouse) for 2 h at room temperature. Sections were washed twice in PBS, mounted on microscope slides, and coverslipped with Vectashield containing DAPI counterstain. Imaging was performed on a Leica Stellaris confocal microscope.

### Statistical analysis

Behavioral and neural activity data were analyzed with a combination of linear mixed effects ANOVAs, planned t tests, and Pearson correlations. Post-hoc comparisons were completed with Bonferroni corrections. Summary figures represent all individuals or averages of subjects with mean ± s.e.m. Statistical significance was set at p<0.05. Photometry and video data were analyzed in MATLAB. Visualization and statistical analyses were performed using Graphpad Prism 10.0.

## Supplemental Materials

**Supplemental Fig 1.**
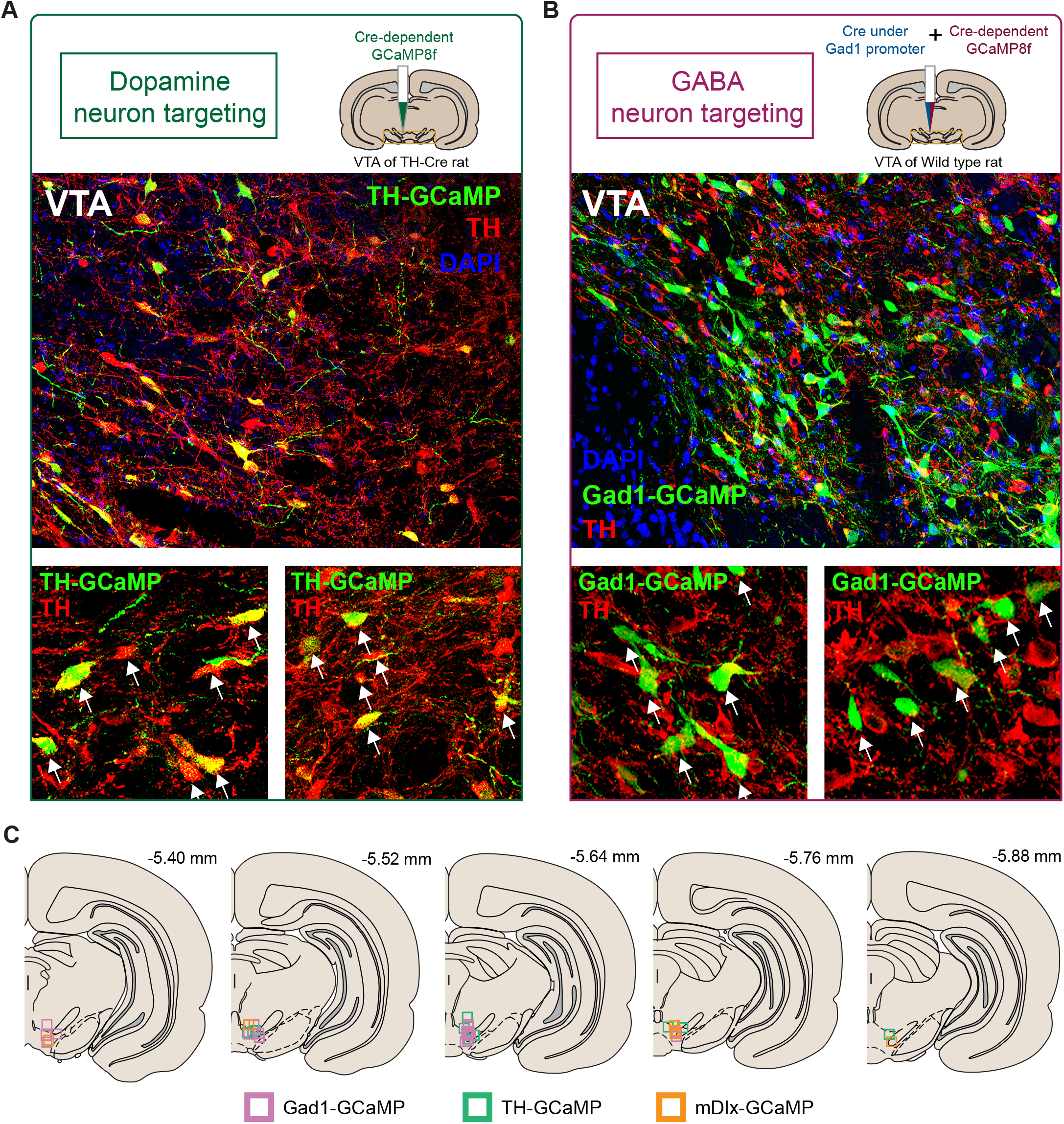
Histology and fiber placements in the VTA for GABA and DA targeted GCaMP recordings. **A)** Dopamine/TH+ neurons were targeted with a cre-dependent AAV in TH-cre rats, with good specificity. Bottom, zoom in on example brain images. White arrows highlight GCaMP expression in TH+ neurons. **B)** Using a dual-virus approach, GAD1+ neurons were targeted, resulting in minimal expression in TH+ neurons in the VTA. Bottom, zoom in on example brain images. White arrows highlight GCaMP expression in neurons that are not TH+. **C)** Optic fiber placements for rats in recording experiments. Pink squares represent GAD1-targeted rats. Green squares represent TH-targeted rats. Orange squares represent mDlx-targeted rats.

**Supplemental Fig 2.**
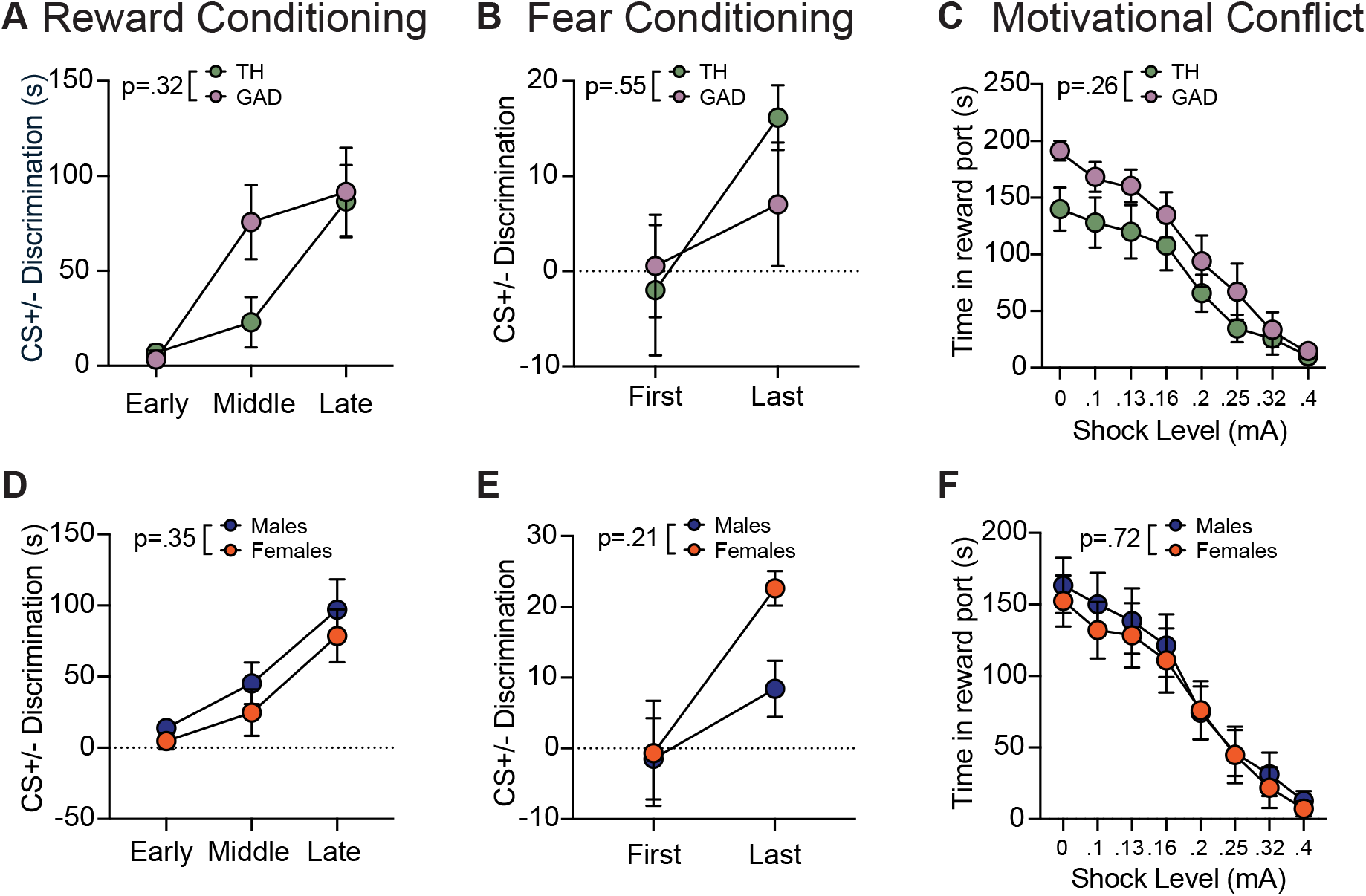
Conditioned behavior did not significantly differ across cohorts. **A)** Gad and TH animals achieved the same level of cue discrimination during reward conditioning as measured by the difference in reward port duration between the CS+ and the CS-, indicating comparable learning. **B)** Gad and TH animals achieved the same level of discrimination during fear conditioning as measured by the difference in percent trials with freezing between the CS+ and the CS-. **C)** Gad and TH animals similarly decreased their cued responding during motivational conflict as shock intensity escalated, as measured by seconds in the reward port. **D)** Male and female animals achieved the same level of discrimination during reward conditioning as measured by the difference in reward port duration between the CS+ and the CS-. **E)** Male and female animals achieved the same level of discrimination during fear conditioning as measured by the difference in percent trials with freezing between the CS+ and the CS-. **F)** Male and female animals decreased responding similarly during motivational conflict as shock intensity escalated, as measured by seconds in the reward port.

**Supplemental Fig 3.**
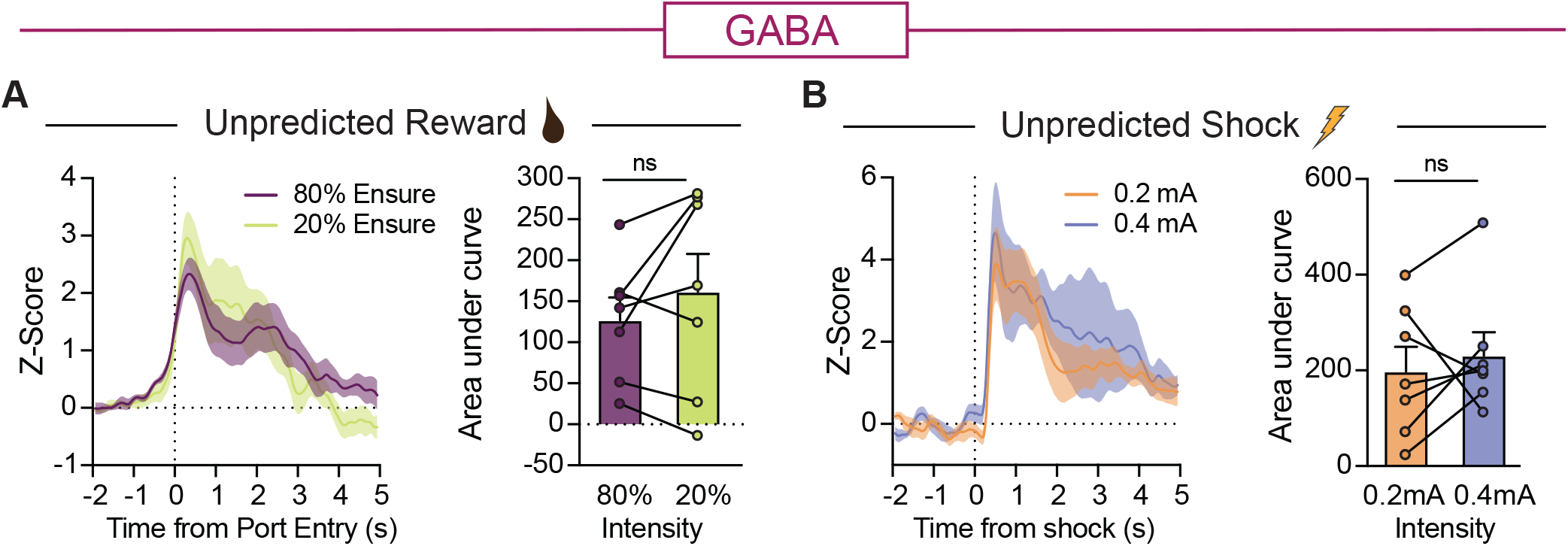
VTA GABA neuron activity does not reliably scale with stimulus intensity. **A)** Reward consumption evoked similar VTA GABA neuron activity regardless of concentration as measured by AUC (paired t test, t(6)=1.18, p=.281). **B)** Footshock evoked similar VTA GABA neuron activity regardless of electrical charge as measured by AUC (paired t test, t(6)=0.65, p=.542).

**Supplemental Fig 4.**
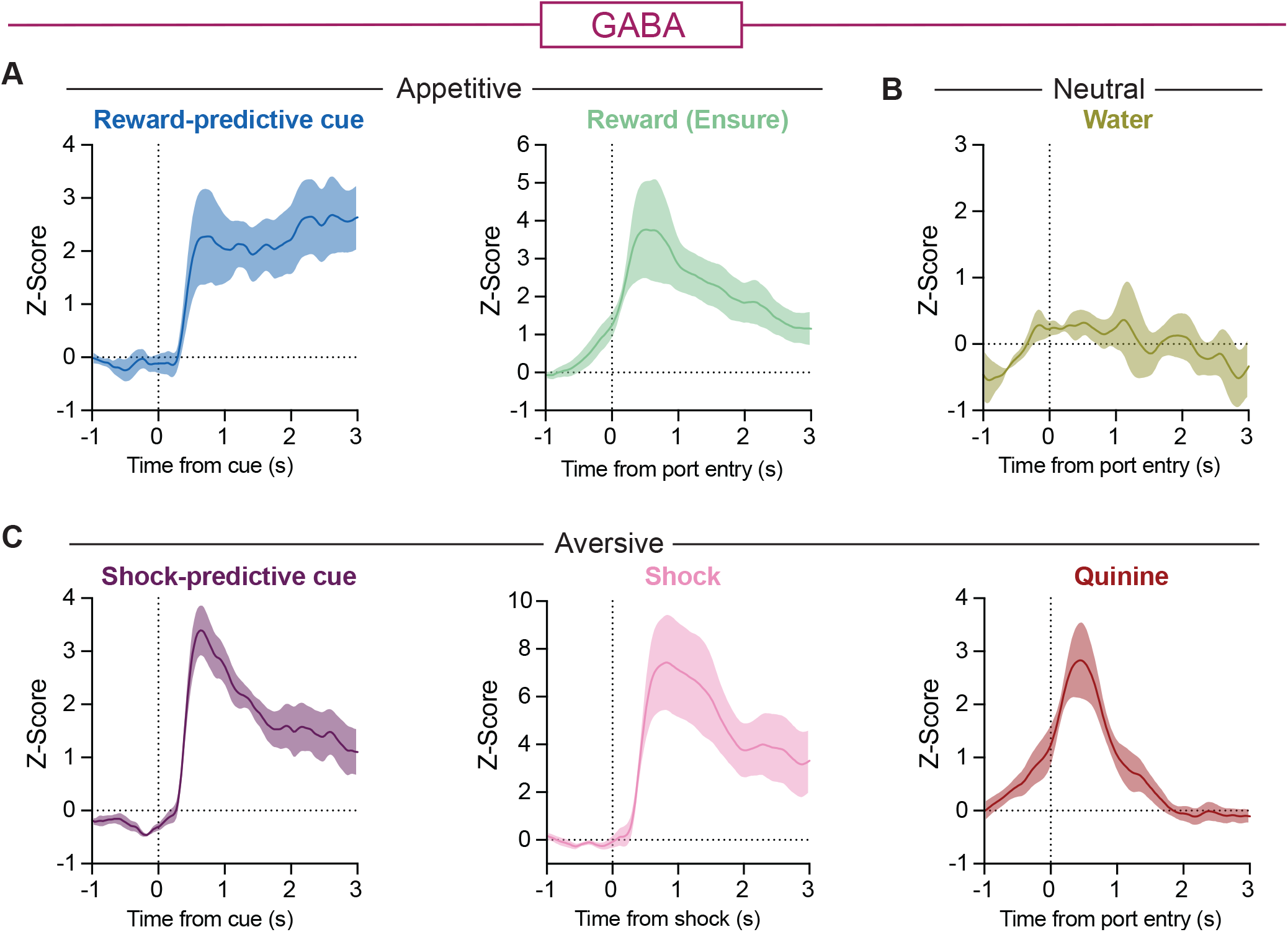
VTA mDlx GABA neurons are activated by emotionally salient stimuli of different valences and modalities. **A)** GCaMP was targeted to another VTA GABAergic population, mDlx+ neurons, followed by photometry recordings during and reward and fear conditioning (n=6). A) Reward and reward-predictive cues consistently activated mDlx+ GABA VTA neurons. **B)** Water consumption, in contrast, did not engage mDlx+ neurons. **C)** Shock and shock-predictive cues consistently activated mDlx+ neurons, and this population was also activated by bitter quinine drinking.

**Supplemental Fig 5.**
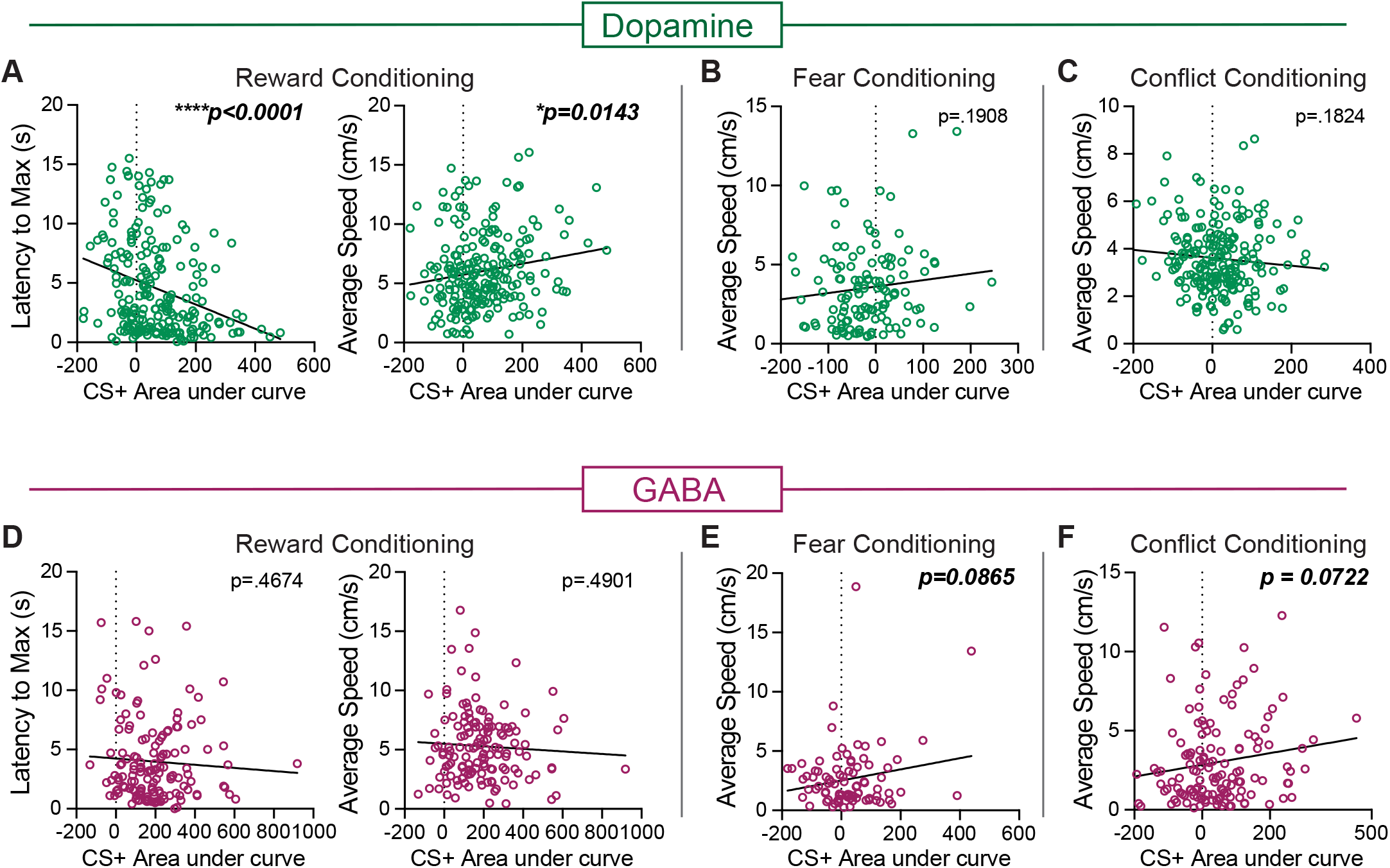
Cue-evoked VTA dopamine and GABA neuron activity differentially predicts movement vigor across appetitive and aversive learning contexts. **A-F)** Pearson correlations between the peak cue-evoked signal and speed for reward conditioning, fear conditioning, and conflict conditioning. **A)** Cue-evoked dopamine neuron activity was negatively correlated with the latency to reach maximum speed and positively correlated with average speed for reward conditioning, but was not correlated with speed for **B)** fear or **C)** conflict conditioning. **D)** Cue-evoked GABA neuron activity was not correlated with speed for reward conditioning, but there was a trend for positive correlations with speed for **E)** fear and **F)** conflict conditioning.

